# A split ALFA tag-nanobody system for protein localization and proximity proteomics in mycobacteria

**DOI:** 10.1101/2025.03.22.644702

**Authors:** Allison Fay, Andrew P. Kurland, Zhuoning Li, Mara Monetti, Jeffrey R. Johnson, Michael S. Glickman

## Abstract

Tuberculosis remains a globally significant infection and new insights into the biology of *Mycobacterium tuberculosis* are badly needed. Discovery of protein localization and protein complex composition are powerful approaches to determine protein function, but have not been widely applied in mycobacteria, in part due to technical barriers. Here we develop a multi-functional system that utilizes the ALFA tag and functional protein fusions to an anti-ALFA nanobody (NBALFA) to target proteins in fast and slow growing mycobacteria. Insertion of the ALFA epitope tag on the target protein, coupled with conditional expression of the NBALFA fused to a fluorescent protein faithfully recapitulates cytosolic and membrane protein localization by fluorescent microscopy in living cells. Targeted NBALFA can relocalize an ALFA tagged protein to inclusion bodies or the cytoplasmic membrane, demonstrating enforced protein localization. Finally, conditional expression of the NBALFA fused to TurboID for proximity proteomics allowed identification of known partner proteins of the RNA polymerase complex and the PKS13 mycolic acid biosynthesis protein. We conclude that the split ALFA tag-nanobody system is a flexible platform for discovery protein biology in mycobacteria.

## Introduction

*Mycobacterium tuberculosis* remains a high burden global health problem, infecting over 10.8 million people and causing 1.25 million deaths in 2023. There were about 400,000 estimated MDR/RR-TB cases in 2023 limiting the available drugs (1). Although our knowledge of the biology of *M. tuberculosis* is expanding, we still lack a complete understanding of the key processes in the mycobacterial cell, including cell wall biosynthesis, transcription, and DNA replication. Understanding the biology of these basic cellular systems is critical for the development of new antimicrobial therapies by elucidating new drug targets or understanding how current therapies affect cell function.

Decades of development of genetic tools for mycobacteria have powered directed and genome wide discovery approaches, including systems for genetic deletion, integration, and point mutagenesis (2–4), CRISPRi technology for high throughput library screening and targeted gene knockdown (5, 6), CRISPR gene targeting for genetic deletions (7, 8), and approaches such as ORBIT which utilize phage machinery to make multiple deletions and chromosomal integrations (9). Although these advances in mycobacterial genetics have dramatically accelerated discovery science for both pathogenic and nonpathogenic mycobacteria, proteomic approaches to understand mycobacterial proteins in their native cellular context have lagged. Protein fusions to a variety of functional proteins, including epitope tags, fluorescent proteins, and chemically modifiable functional domains allow for interrogation of protein spatial positioning, environment, and binding partners. Localization via fluorescent protein fusions have elucidated a vast array of spatial patterning within mycobacterial cells including, but not limited to asymmetric polar growth elucidated by GFP-Wag31 localization (10), proteostasis machinery utilizing the tagged chaperone DnaK-mCitrine and disaggregase ClpB-mCitrine (11), and replication and architecture of the chromosome with tagged nucleoid-associated proteins, SSB-GFP and HupB-GFP (12, 13). High-throughput fusion libraries have also been developed to systematically assess protein patterning (14, 15). However, direct protein fusions to large partner proteins can introduce artifacts and impair protein function. For live cell localization, the most common fusion protein, GFP, though generally well tolerated, can impact folding, stability, and function due to its large size (16). GFP fusions can alter protein localization in live cells (17) and generally must be generated at the N- or C-terminus of the target.

Proximity proteomics is a powerful technique to investigate protein function by capturing both stable and transient protein interactions (18). The most commonly used systems fuse APEX/APEX2 (19, 20) or the BirA-derivatives, BioID/TurboID (21, 22), to a target protein of interest, thereby directing biotinylation to partner proteins in proximity. Recently, APEX-based proximity labeling was used in mycobacteria to identify the cytoplasmic and periplasmic compartment proteomes in Mtb (23, 24) and a BioID fusion was used to interrogate HbhA in *M. smegmatis* (25), but otherwise proximity proteomics has not been widely deployed in mycobacteria.

The ALFA tag is a rationally designed epitope tag that is small, balanced, alpha-helical, minimally reactive, and amenable to placement at the termini or internally in a target protein (26). Critically, a cognate nanobody (NBALFA), a single chain camelid antibody with picomolar binding affinity, was generated as a partner reagent for the ALFA tag (26). Because nanobodies efficiently fold and bind their target sequence within the prokaryotic cell cytoplasm (27), the ALFA-NBALFA pair would potentially provide a flexible system in which a single ALFA tagged target could be functionally targeted in live mycobacterial cells by fusing partner proteins to the NBALFA. This manuscript explores this idea and presents a multifunctional NBALFA/ALFA tag system for protein studies in fast and slow growing mycobacteria.

## Results

### Nanobody based protein localization in *M. smegmatis*

The design of the split ALFA system is presented in Figure 1. To test the utility of this approach, we generated chromosomal C-terminal ALFA tagged proteins by integrating the ALFA tag at the 3’ end of the endogenous *rpoC* or *mmpL3* genes in *M. smegmatis*. These proteins are well characterized in mycobacteria with clear patterns of localization: RpoC to the nucleoid and MmpL3 to cell poles, (28–30). *M. smegmatis* MmpL3-ALFA and RpoC-ALFA were viable and had similar replication times in culture, indicating the ALFA tagged versions of both proteins were functional given that both genes are essential for viability. We verified expression of each ALFA tagged protein at the predicted size via immunoblot using NBALFA (Figure 2A). We then introduced plasmids encoding GFP translationally fused at the C-terminus to NBALFA with flexible linker (GGGSGGG) expressed from *ftsZ* (P_ftsZ_), *hsp60* (P_hsp60_), or the Tet-inducible promoters into wildtype *M. smegmatis*, RpoC-ALFA, or MmpL3-ALFA. The protein levels of GFP-NBALFA correlated with expected promoter strength with P_ftsZ_ showing very little full length fusion protein compared to the P_hsp60_ driven construct (Figure 2B). Tet-inducible GFP-NBALFA was undetectable without inducer, but accumulated at its predicted size with anhydrotetracycline (ATc) induction (Figure 2B).

**Figure 1.**
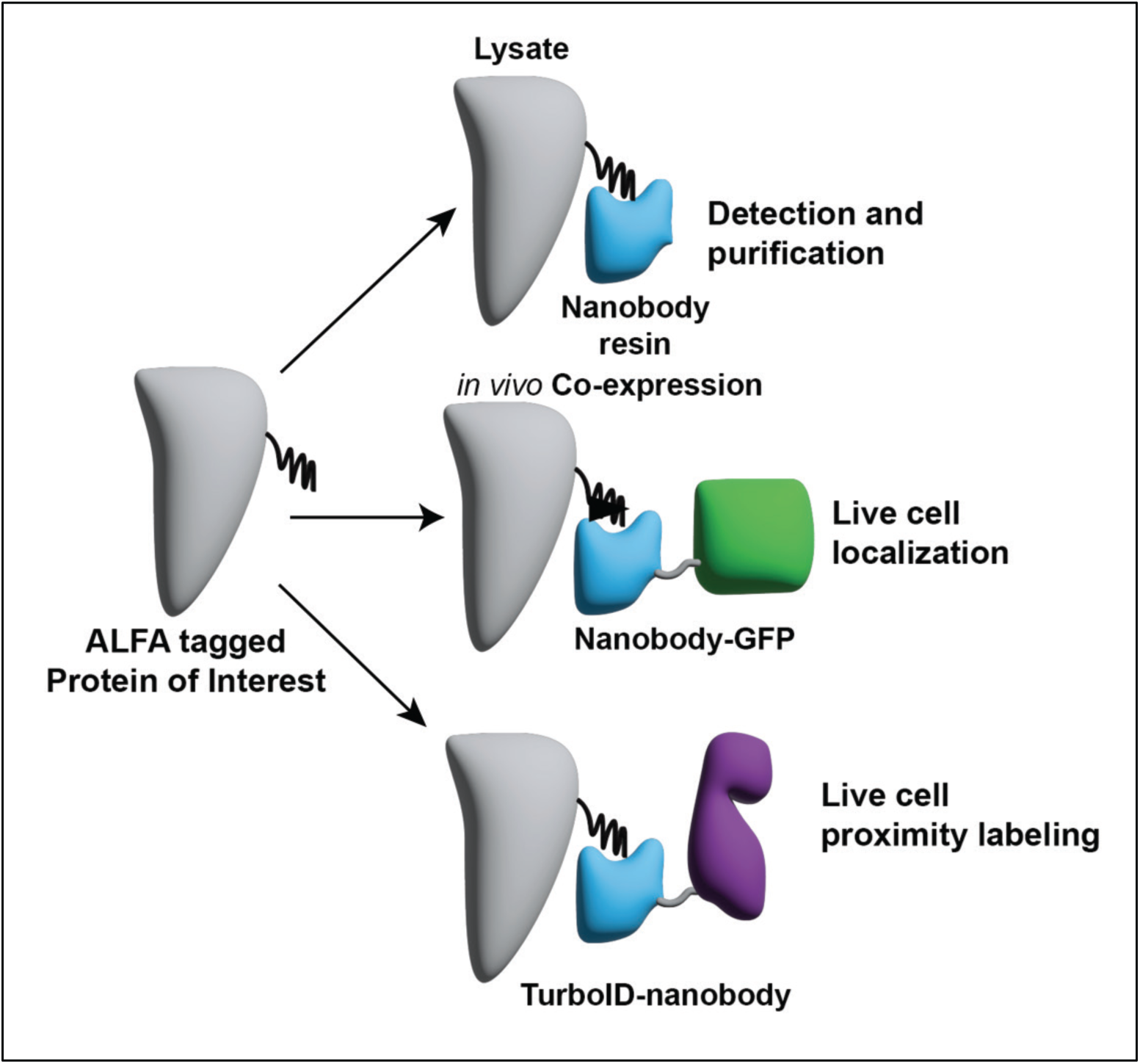
Design of a split ALFA tag nanobody system for detection/purification, localization, and proximity labeling in mycobacteria.

**Figure 2.**
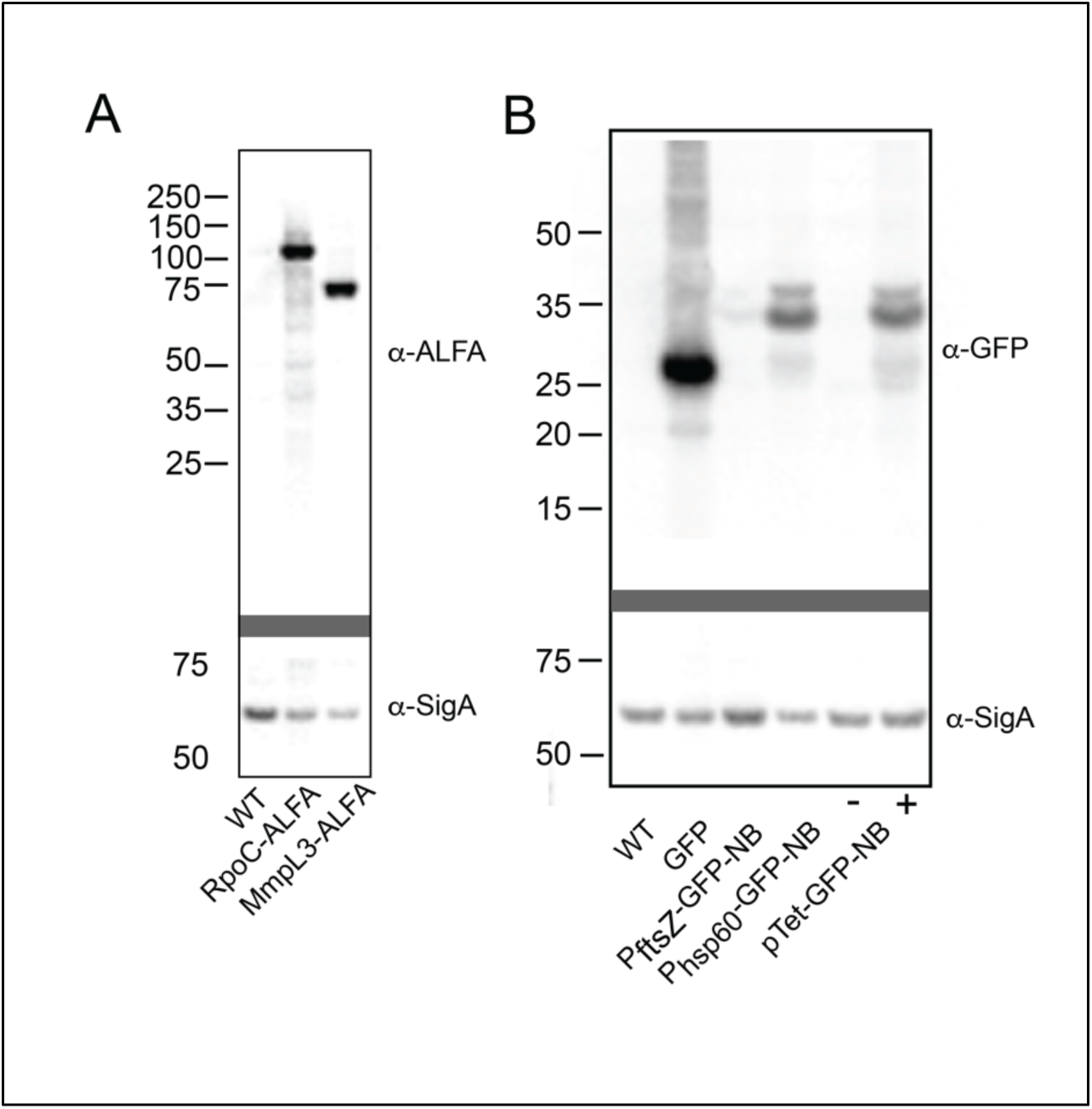
Chromosomal ALFA tagged MmpL3 and RpoC. **(A)** Immunoblots of lysates from MC2155 (WT), MGM7103 (encoding RpoC-ALFA), and MGM7059 (encoding MmpL3-ALFA). The top blot is probed using NBALFA and anti-VHH-HRP. The bottom blot stripped and probed with anti-SigA and anti-rabbit Ig-HRP as a loading control. **(B)** Immunoblots of lysates from MC2155, MGM6970 (expressing GFP), MGM7087 (expressing GFP-NBALFA from the P_ftsZ1_ promoter), MGM7089 (expressing GFP-NBALFA from the P_hsp60_ promoter), MGM7047 (expressing TetON GFP-NBALFA, no ATc), and MGM7047 (encoding TetON GFP-NBALFA, + ATc-50ng/ml). Top blot probed with anti-GFP and anti-rabbit Ig-HRP. Bottom blot stripped and probed with anti-SigA and anti-rabbit Ig-HRP.

To determine if NBALFA can localize to ALFA tagged proteins in living cells, we imaged GFP-NBALFA by fluorescent microscopy in the absence or presence of an ALFA tagged target protein. Without an ALFA tagged protein, fluorescent signal from GFP-NBALFA was diffusely distributed and indistinguishable from free GFP (Figure 3A), with fluorescence signal intensity correlating to protein levels seen by immunoblot. We next assessed whether GFP-NBALFA would localize to ALFA tagged proteins *in vivo*. Mycobacterial RpoC-GFP was previously reported to co-localize to the nucleoid in logarithmically growing cells (28) and an RpoC-mCitrine fusion protein similarly colocalized with the irregular puncta of the nucleoid visualized by Hoechst staining (Figure 3B top panels). In RpoC-ALFA expressing cells, the nucleoid localization pattern was weakly visible in cells co-expressing the GFP-NBALFA under P_ftsZ_ and was easily visible when GFP-NBALFA was constitutively or inducibly expressed at higher levels (Figure 3B). The pattern of GFP-NBALFA localizing to RpoC-ALFA was indistinguishable from the direct RpoC-mCitrine fusion and contrasted with the diffuse pattern seen in cells not expressing an ALFA-tagged protein. We repeated these experiments with MmpL3-GFP and MmpL3-ALFA fusions. An MmpL3-GFP translational fusion accumulated at the poles and septa as previously described (29, 30). Similarly, *M. smegmatis* cells expressing MmpL3-ALFA and high levels of GFP-NBALFA (P_hsp60_ or Tet-induced) also showed a polar localization pattern that was similar to the direct MmpL3-GFP fusion (Figure 3C). These results indicate that GFP-NBALFA will localize to cytoplasmic or membrane proteins when co-expressed with an ALFA tagged target.

**Figure 3.**
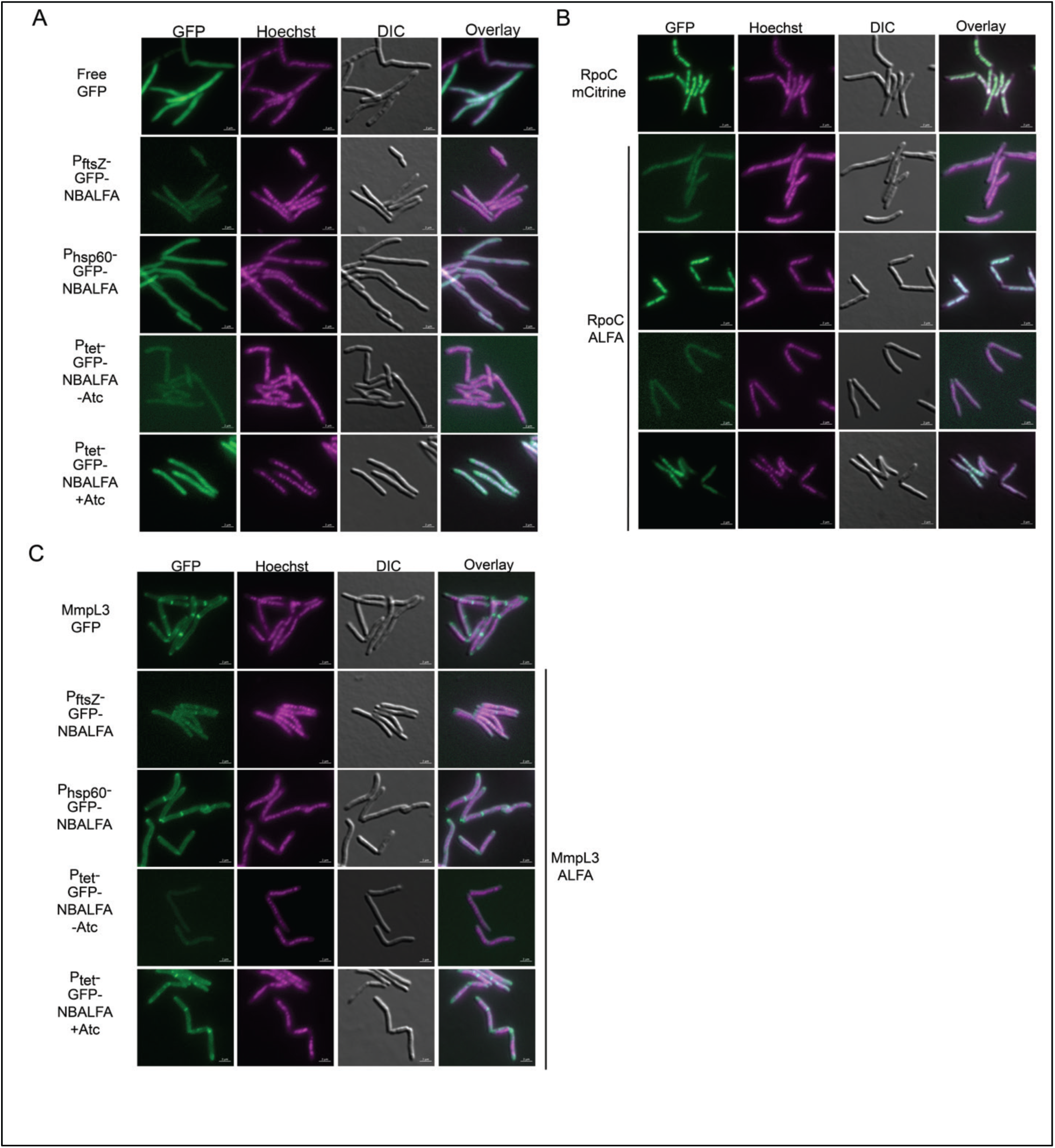
Detection of protein localization in live mycobacterial cells using split NBALFA. **(A)** GFP-NBALFA mimics the distribution of free GFP in the absence of an ALFA tagged target protein. MGM6970 (expressing free GFP), MGM7087 (expressing GFP-NBALFA from P_ftsZ1_), MGM7089 (expressing GFP-NBALFA from P_hsp60_), MGM7047 (expressing TetON GFP-NBALFA, no ATc), MGM7047 (encoding TetON GFP-NBALFA, + ATc-50ng/ml). **(B)** NBALFA localizes to RpoC in living cells. MGM6026 (expressing an RpoC-mCitrine protein fusion), MGM7107 (expressing RpoC-ALFA and GFP-NBALFA under P_ftsZ1_), MGM7108 expressing RpoC-ALFA and GFP-NBALFA under P_hsp60_), MGM7105 (expressing RpoC-ALFA and TetON GFP-NBALFA, no ATc), MGM7105 (expressing RpoC-ALFA and TetON GFP-NBALFA, + ATc-50ng/ml). **(C)** NBALFA localizes to MmpL3 in living cells. MGM6464 (expressing a MmpL3-GFP fusion), MGM7099 (expressing MmpL3-ALFA and GFP-NBALFA under P_ftsZ1_), MGM7101 (encoding MmpL3-ALFA and GFP-NBALFA under P_hsp60_), MGM7081 (expressing MmpL3-ALFA and TetON GFP-NBALFA, no ATc), MGM7081 (expressing MmpL3-ALFA and TetON GFP-NBALFA, + ATc-50ng/ml). All images contain 2uM scale bar and show GFP (imaged 500ms, 70% Colibri LED), Hoechst staining (DAPI, 500ms external lamp), DIC, and overlay.

### Protein Relocalization using NBALFA

We next tested whether we could re-target an ALFA tagged protein to distinct subcellular locations using NBALFA targeted to the membrane or a protein inclusion body. For inclusion body targeting, we added the ELK16 aggregation sequence, which forms aggregates and inclusion bodies *in vivo* (31), at the N-terminus of GFP-NBALFA to create ALFA tagged fluorescent insoluble aggregates. When viewed by fluorescent microcopy, unlike the diffuse cytoplasmic signal of GFP-NBALFA (Figure 3A), ELK16-GFP-NBALFA formed multiple foci in the cell that occasionally coalesced into a large, polar inclusion bodies (Figure 4A). To test whether ALFA tagged proteins would re-localize to these inclusion bodies, we coexpressed the Tet-inducible ELK16-GFP-NBALFA with mCherry-ALFA or mCherry-HA. Upon induction of ELK16-GFP-NBALFA and aggregate formation, mCherry-ALFA colocalized with GFP-NBALFA puncta (Figure 4A, top). This relocalization was specific to the ALFA tag, as mCherry-HA remained in a diffuse pattern when coexpressed with ELK16-GFP-NBALFA aggregates (Figure 4A, bottom).

**Figure 4.**
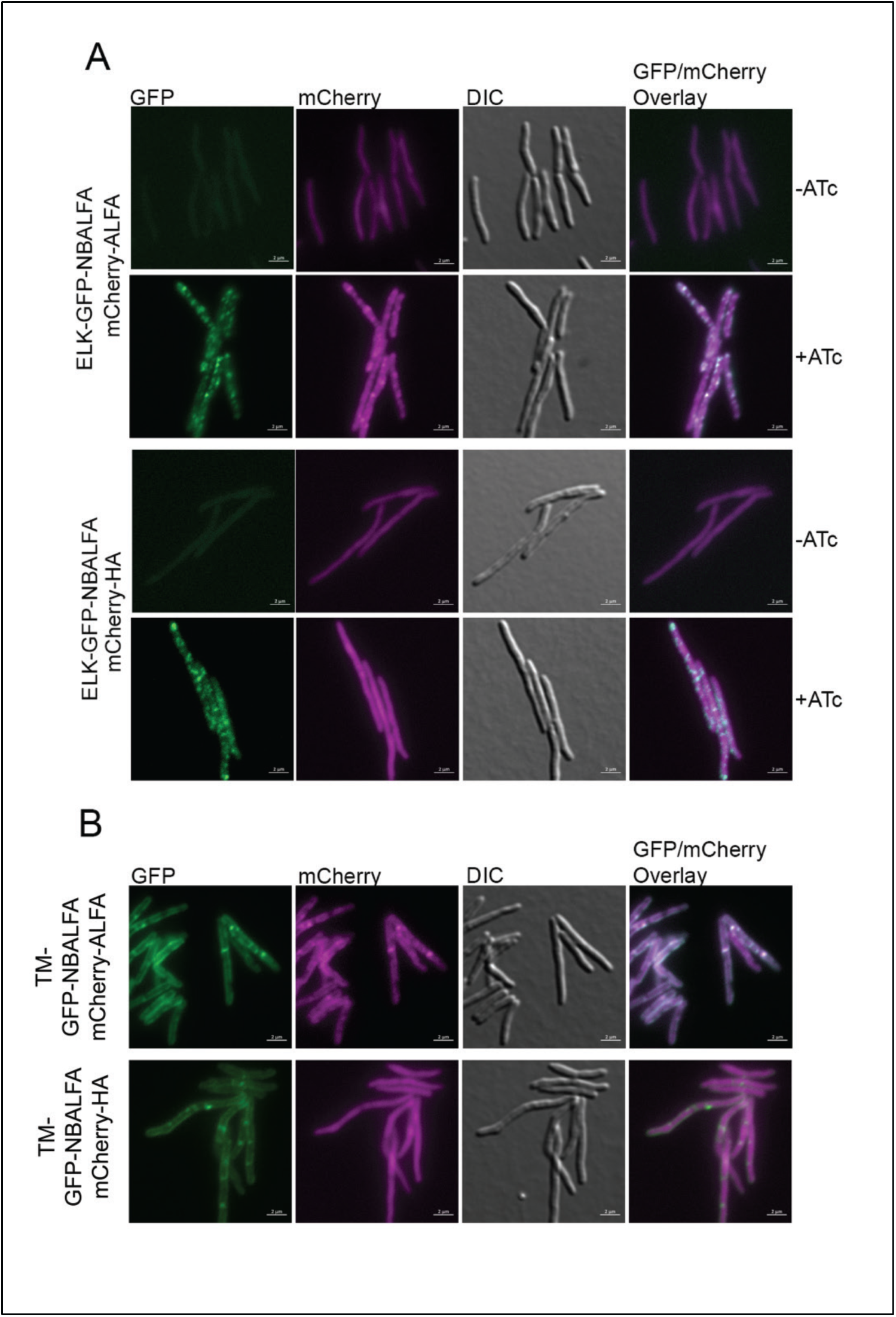
Enforced protein relocalization using split NBALFA. **(A)** Relocalization to inclusion bodies. Top section: AFM1996 (mCherry-ALFA, TetON ELK16-GFP-NBALFA) without (top) and with (bottom) ATc showing GFP inclusion body formation and colocalization of mCherry-ALFA to inclusions. Bottom section, same as (A), but with a control mCherry-HA expression strain, AFM1995 (mCherry-HA, TetON ELK16-GFP-NBALFA) without (top) and with (bottom) ATc to induce GFP inclusion bodies. **(B)** Relocalization to the membrane. mCherry-ALFA, or control mCherry-HA, with coexpression of an inducible membrane targeted MalF_(TM1,2)_-GFP-NBALFA. Top section: MGM7879 (mCherry-ALFA, TetON MalF_(TM1,2)_-GFP-NBALFA) with ATc. Bottom section: MGM7985 (mCherry-HA, TetON MalF_(TM1,2)_-GFP-NBALFA) with ATc. All images contain 2uM scale bar and show GFP (imaged 500ms, 70% Colibri LED), mCherry (imaged 500ms, 70% Colibri LED), DIC, and GFP/mCherry overlay.

To test whether NBALFA could relocalize an ALFA tagged protein to the membrane, we utilized the first two transmembrane domains of *E. coli* MalF, which we previously used to mark *M. smegmatis* membranes (11, 29). MalF_(1,2)_-GFP-NBALFA localized in a membrane and septal pattern in *M. smegmatis* (Figure 4B) and coexpression of mCherry-ALFA relocalized cherry signal to the membrane (Figure 4B, top), whereas mCherry-HA remained in a diffuse cytoplasmic pattern (Figure 4B, bottom). These experiments demonstrate that the split ALFA-NB system can be used to relocalize proteins in living mycobacterial cells.

### Split ALFA nanobody based proximity proteomics in mycobacteria

After confirming the NB based localization to ALFA tagged proteins in living *M. smegmatis*, we adapted biotin-based proximity labeling to the split ALFA system. We chose the promiscuous biotin-ligating enzyme, TurboID, which can label available lysines with biotin (21). We generated a conditional expression vector encoding TurboID translationally fused to NBALFA (Figure 1). When expressed in *M. smegmatis* RpoC-ALFA or MmpL3-ALFA, we were unable to detect TurboID-NBALFA in uninduced cell lysates (Figure 5A). After induction for 3 hours with ATc, the TurboID-NBALFA fusion protein was visible at the expected size when probed with an antibody to BirA, (Figure 5A).

**Figure 5.**
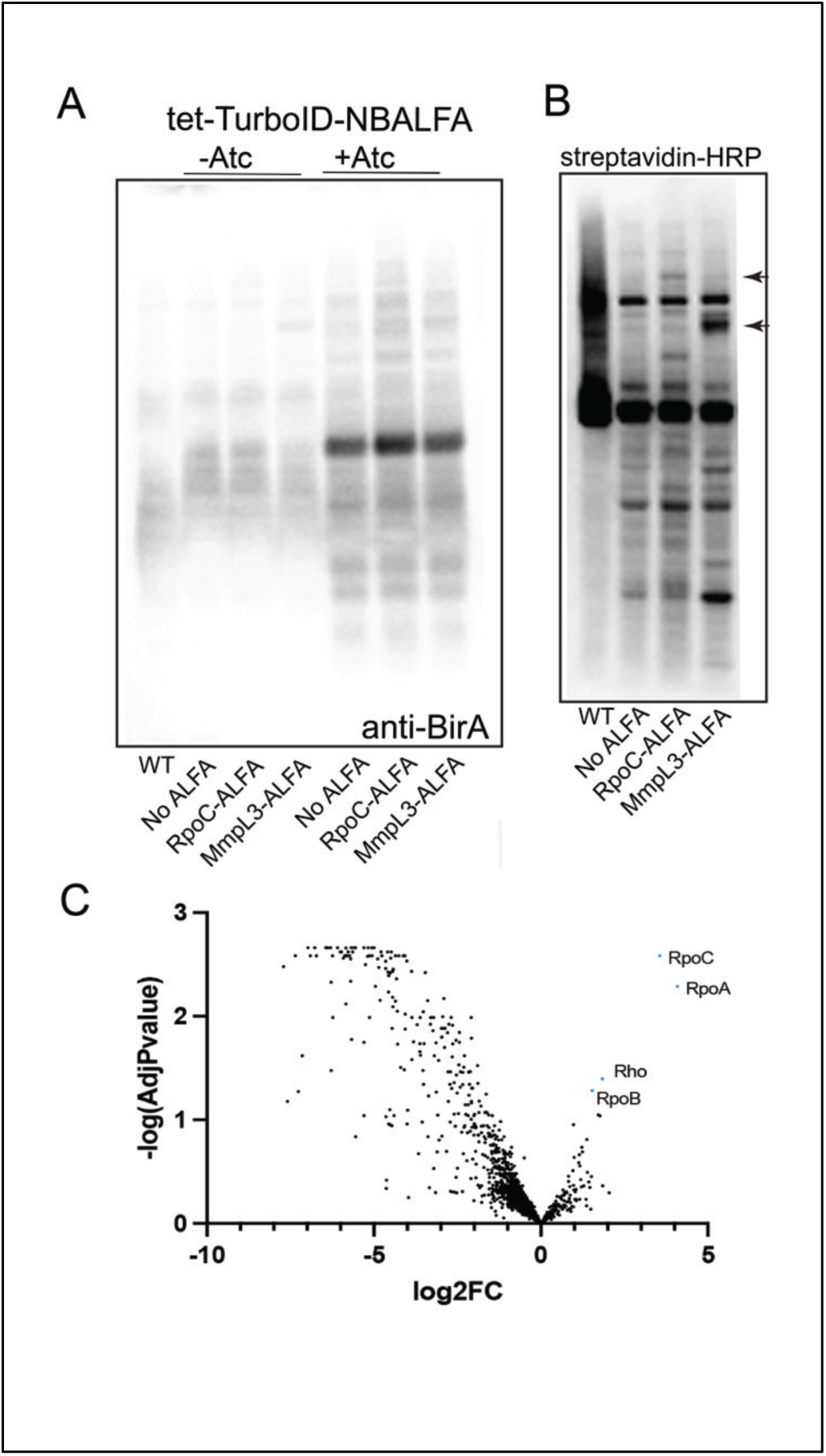
Proximity labeling in *M. smegmatis* expressing ALFA tagged RpoC or MmpL3 with a TurboID-NBALFA fusion. **(A)** Immunoblots of lysates from WT *M. smegmatis*, MGM7085 (no ALFA), MGM7105 (encoding RpoC-ALFA), MGM7081 (encoding MmpL3-ALFA), all with a tet inducible TurboID-NBALFA. Lanes 2-4 are uninduced and lanes 5-7 were induced with ATc-50ng/ml for 3 hours. The blot was probed with anti-BirA then anti-rabbit Ig-HRP. **(B)** Streptavidin-HRP probed blot to detect biotinylated proteins in WT, no ALFA, RpoC-ALFA and MmpL3-ALFA, all expressing TurboID-NBALFA after biotin labeling. Black arrowheads indicate biotinylated proteins at the predicted size of RpoC-ALFA (lane 3) or MmpL3-ALFA (lane 4) **(C)** Volcano plot of RpoC-ALFA compared to No ALFA proximity labeling. Blue points indicate Log2FC>1 and Adjusted P-value <0.05. Significant blue points labeled with protein names, including RpoC bait protein.

To determine if the TurboID-NBALFA fusion protein could catalyze biotinylation of an ALFA tagged target, we developed a labeling protocol that would limit non-specific labeling by employing transient expression of the TurboID-NBALFA fusion to allow localization to the ALFA tagged target before initiation of labeling with biotin addition. However, mycobacterial cells synthesize biotin (32, 33) and produce many biotinylated proteins (8, 34–36), indicating that TurboID catalyzed biotinylation might proceed without biotin addition. To optimize conditions for labeling, we measured TurboID-NBALFA dependent biotinylation in different media conditions. Wild type *M. smegmatis* without TurboID has several dominant biotinylated proteins detected by streptavidin-HRP (Figure S1). *M. smegmatis* expressing tet inducible TurboID-NBALFA in biotin free Sauton’s media with addition of inducer were indistinguishable from WT cells, indicating that the endogenous biotin pool is not available for TurboID-NBALFA catalyzed biotinylation, at least at this level of detection. However, expression of TurboID-NBALFA in LB media resulted in detectable additional biotinylated proteins, and addition of biotin to Sauton’s media resulted in the highest level of additional biotinylation (Figure S1). These results indicate that there is background endogenous biotinylation by TurboID in mycobacteria, but that biotin free media and exogenous biotin addition can temporally restrict this background. However, these results also emphasize the importance of the appropriate negative ALFA tagged control proteins to detect specific biotinylation signals above both endogenously biotinylated proteins and those nonspecifically biotinylated by TurboID.

To test the system in cells expressing an ALFA tagged target protein, we used TurboID-NBALFA expressing cells carrying either RpoC-ALFA and MmpL3-ALFA. We observed biotinylated proteins corresponding to the predicted sizes of the bait proteins, suggesting that the TurboID-NBALFA was targeted to the ALFA tagged bait protein and subsequently labeled the bait protein with biotin (Figure 5B). In addition, there were additional new biotinylated proteins in the lysates each of the ALFA expressing cells compared to no ALFA control cells (Figure 5B), suggesting proximity labeling of additional non-bait proteins above background. To identify these additional biotinylated proteins, we isolated biotinylated proteins from RpoC-ALFA and no ALFA control cells with streptavidin-agarose and quantitated peptides by mass spectrometry. We identified the bait protein, RpoC, as a top hit enriched in the RpoC-ALFA strain when compared to the no ALFA tag control strain (Figure 5C and Table S2). Furthermore, we identified the core RNAP subunits RpoA (RNAPα) and RpoB (RNAPβ) as well as the transcription termination protein, Rho as RpoC proximity hits (Figure 5C and Table S2).

### Proximity labeling in *M. tuberculosis*

To test proximity labeling using TurboID-NBALFA in *M. tuberculosis*, we targeted PKS13, a large (186 kDa) multimodular protein essential for mycobacterial viability (37–39). PKS13 is essential for mycolic acid biosynthesis and condenses the alpha branch fatty acids with the meromycolate chain in the final steps of mycolic acid synthesis (37). Due to its distinct structural domains, PKS13 is an ideal candidate to test the potential for domain specific proximity labeling by placing the ALFA tag at distinct sites in the protein. The ALFA tag’s small size and balanced α-helical structure was reported to be tolerated at termini and at internal sites (26). We placed the ALFA tag at the N-terminus, C-terminus, and two internal sites (552AA or 1357AA) in PKS13. The internal sites were chosen for their predicted disordered structure between known functional domains (39). To test the ability of different PKS13 ALFA tagged alleles to complement the essential function of PKS13, we generated *M. bovis* BCG Δ*BCG_3862c* in a merodiploid strain with a second copy of *rv3800c* (encoding *M. tuberculosis* PKS13) at the *attB* site containing a nourseothricin resistance marker. L5 integrase catalyzed allelic exchange with each ALFA-tagged PKS13 encoding plasmid conferring streptomycin resistance confirmed that all four PKS13-ALFA alleles were functional (Figure S2).

To test PKS13 proximity labeling in *M. tuberculosis*, a constitutively expressed second copy of *rv3800c*, the gene encoding PKS13, was integrated into the *M. tuberculosis* chromosome. All *pks13* encoding alleles with distinct ALFA tag positions expressed an ALFA tagged protein at the predicted size of PKS13, with subtle differences in PKS13 levels with different tag positions, as compared to the DnaK loading control (Figure 6A). Coexpression of – TurboID-NBALFA and subsequent biotin labeling revealed a high molecular weight streptavidin reactive protein in all triplicate samples consistent with the expected size of PKS13, which was not visible in the no ALFA control (Figure 6B). To identify biotinylated proteins dependent on the ALFA-TurboID-NBALFA interaction, we purified biotinylated proteins from each of the labeled lysates on streptavidin resin and quantitated peptides by mass spectrometry. PKS13 was the top enriched protein identified by mass spectrometry in all four ALFA tag positions, indicating proper targeting and labeling by TurboID-NBALFA, independent of tag position (Figure 6C, Table S3). The PKS13 proximity interactome included multiple proteins with known functions in mycolic acid biosynthesis including early Fas1 associated factors (AcpM, AccD6), FasII components (InhA, HadB), mycolic acid methyltransferases that modify the meromycolate chain (MmaA3, MmaA4), and proteins involved in TMM transport (CmrA) (Figure 6C, S3 and Table S4). Additional proteins with relevance to the mycolic acid pathway were also detected (Figure 6C, Table S3), including proteins in the TAG lipid synthesis and lipid degradation pathways (TGS1 and FadE23) whose products can be used for mycolic acid synthesis, (40–42). We also identified protein chaperones, including DnaK, which we previously implicated as required for large multimodular protein stability in *M. smegmatis* (11). To determine if distinct ALFA tag positions within PKS13 led to distinct biotinylation patterns, we compared the proximity interactome of PKS13 alleles with different ALFA tagged positions. This analysis revealed a core PKS13 interactome shared in all four ALFA tagged PKS13 positions (Figure 6D, Table S5), but also revealed proximity hits unique to each tag position, particularly for the 1352AA tag position, indicating that ALFA tag placement can reveal distinct protein interactors in large multidomain proteins.

**Figure 6.**
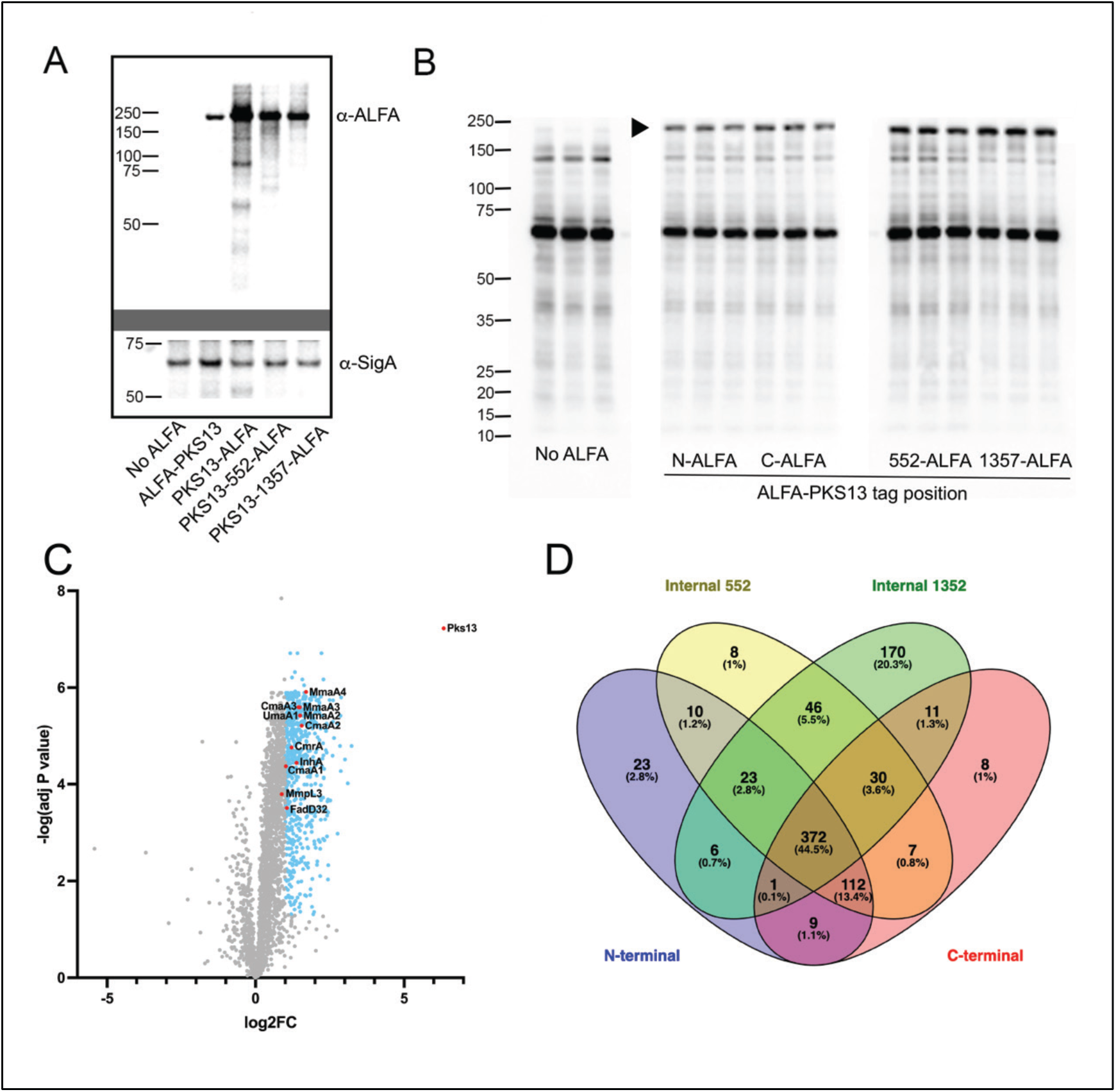
Proximity labeling of *M. tuberculosis* PKS13. **(A)** Immunoblots of strains expressing ALFA tagged PKS13 proteins: no ALFA tag, N-terminal ALFA tagged PKS13, C-terminal ALFA tagged PKS13, internal ALFA tag (552AA) PKS13, and Internal ALFA tag (1357AA) PKS13. Top probed using NBALFA then anti-VHH-HRP. Bottom blot reprobed with anti-SigA then anti-rabbit Ig-HRP. **(B)** Streptavidin-HRP detection of biotinylated proteins in triplicate proximity labeling experiments in *M. tuberculosis* no ALFA control, N-terminal ALFA tagged PKS13, C-terminal ALFA tagged PKS13, internal ALFA tag (552AA) PKS13, and internal ALFA tag (1357AA) PKS13. Arrowhead indicates a biotinylated protein of approximate bait (PKS13) size. **(C)** Volcano plots of proximity labeling samples N-terminal ALFA-PKS13 compared to No ALFA. Blue points indicate Log2FC>1 and Adjusted P-value <0.05. with red points highlighting proteins with known functions in mycolic acid biosynthesis. **(D)** Venn diagram depicting position specific proximity interactome (Log2FC > 1 and Adjusted P-value < 0.001) of PKS13 (also see Table S5 for specific protein lists).

## Discussion

We have created a flexible split ALFA tag-ALFA nanobody based system for protein-based discovery science in mycobacteria. The core enabling technology of this system is the ALFA tag/NBALFA pair. The high affinity of the NBALFA for the ALFA tag, combined with the ability to express the single chain in living bacterial cells, enables our approach of using the NB as a delivery vehicle to localize distinct functional protein partners to an ALFA tagged protein of interest. This design gives tremendous flexibility as it allows a single bacterial strain bearing an ALFA tagged protein of interest to be used for protein localization, affinity purification, and proximity proteomics, simply by changing the fusion partner of the expressed NBALFA. The split design of the system may offer several advantages in addition to this flexibility. The small size and optimized structural design of the ALFA tag is presumably less likely to artifactually perturb protein function or localization than a larger direct fusion such as a fluorescent protein or TurboID. The functionalized NB can also be conditionally expressed, allowing temporal control of the imaging or proximity output to coincide with experimental conditions. We demonstrate the utility of the system for visualizing the subcellular distribution of both cytoplasmic and membrane proteins by fluorescent microscopy; for tethering a protein to a specific subcellar compartment; and for proximity proteomics. This latter usage, proximity proteomics, provides a new system to interrogate protein complexes in mycobacteria. Although APEX based proximity proteomics has been applied in mycobacteria for the periplasmic and cytoplasmic compartment, this system is less amendable to cytoplasmic labeling has not been demonstrated for a specific protein target (23, 24, 43). In contrast, the NBALFA system should be useful for cytoplasmic proteins and membrane proteins if targeted to the cytoplasmic domains of those membrane proteins. We also note that the ALFA tag can also be used directly for affinity purification of protein complexes, with or without crosslinking, providing an approach for structural characterization of native protein complexes purified directly from mycobacteria.

We can envision several future uses of this system for discovery biology in mycobacteria. It would be possible with this system to interrogate the proximity interactome of an entire pathway by ALFA tagging a set of proteins with sequential steps in the pathway. Similarly, the dynamic protein interactome of *M. tuberculosis* antibiotic targets could be interrogated during drug treatment or other exogenously applied cellular stress. Similarly, additional functional domains could be fused to NBALFA to enable additional discovery approaches, such as HaloTag for small molecule labeling. We also envision that this approach may be generally applicable to other bacterial or eukaryotic systems in which the flexibility of the split ALFA approach would offer similar advantages.

## Methods

### Growth conditions

*M. smegmatis* strains were cultured in LB with 0.5% glycerol, 0.5% dextrose, and 0.05% Tween 80 (LB_smeg_) or Difco Middlebrook 7H9 supplemented with 0.2% glycerol, 10% ADS (0.5% albumin, 0.085% NaCl, 0.2% dextrose) and 0.05% Tween 80 (7H9). *M. bovis* BCG Pasteur strains were cultured in Difco Middlebrook 7H9 supplemented with 0.2% glycerol, 10% ADS (0.5% albumin, 0.085% NaCl, 0.2% dextrose) and 0.05% Tween 80 (7H9) or on Difco Middlebrook 7H10 agar plates were supplemented with 0.2% glycerol and 10% (wt/vol) Middlebrook oleic acid-albumin-dextrose-catalase (OADC). *M. tuberculosis* Erdman strains were cultured in Difco Middlebrook 7H9 supplemented with 0.2% glycerol and 10% (wt/vol) Middlebrook oleic acid-albumin-dextrose-catalase (OADC) and 0.05% Tween 80 (7H9) or on Difco Middlebrook 7H10 agar plates were supplemented with 0.2% glycerol and 10% (wt/vol) Middlebrook oleic acid-albumin-dextrose-catalase (OADC). Sauton’s media was utilized for biotin-free conditions.

### Bacterial and DNA manipulations

Standard procedures were used to manipulate recombinant DNA and to transform *E. coli*. *M. smegmatis* strains were derivatives of mc2155 (44). *M. bovis* BCG Pasteur and *M. tuberculosis* Erdman were also used. Gene deletions were made by double stranded DNA recombineering (3). All strains used in this study are listed in Table S1. Plasmids including relevant features, and primers are listed in Table S1. *M. smegmatis*, *M. bovis BCG*, and *M. tuberculosis* were transformed by electroporation (2500V, 2.5µF, 1000Ω). Antibiotic concentrations used for selection of mycobacterial strains were as follows: kanamycin 20 µg/ml, hygromycin 50 µg/ml, streptomycin 20 µg/ml, nourseothricin 25 µg/ml.

### Immunoblotting

For protein and epitope tag detection, the following antibodies were used: GFP (Rockland Immunochemicals, Rabbit Anti-GFP polyclonal antibody, 1 mg/ml, 1∶20,000), SigA (purified anti-SigA from rabbit sera, 1∶10,000) (45), TurboID (Invitrogen, BirA Polyclonal Antibody, 1mg/ml, 1:10,000), DnaK (anti-DnaK rabbit poly clonal sera, 1:10,000) (46), ALFA tag (purified NBALFA 1mg/ml, 1:5000 and Genscript, Rabbit anti-VHH HRP conjugated antibody, 1:10,000).

### NBALFA Purification

The sequence encoding the ALFA tag binding nanobody was cloned with a C-terminal 6histidine tag into an IPTG inducible construct, PelB leader-containing derivative of pET21b and expressed and purified as described (47) with the following modifications. A 19-hour induction was done at 22 degrees C in BL21(DE3) cultured in Terrific Broth containing carbenicillin (100ug/ml). Eluates were pooled, concentrated in Amicon Ultra-4 3k, and run on Superdex 200 10/300 size exclusion column using an AKTA purifier in PBS. Nanobody was stored at concentrations of 1-10mg/ml in PBS containing 20% glycerol at −80C.

### Microscopy

All images were acquired using a Zeiss Axio Observer Z1 microscope equipped with Definite focus, Colibri2.0 and Illuminator HXP 120 C light sources, a Hamamatsu ORCA-Flash4.0 CMOS camera, and a Plan-Apochromat 100×/1.4 numerical aperture oil differential inference contrast (DIC) lens objective. Zeiss Zen software was used for acquisition and image export. The following filter sets and light sources were used for imaging: GFP (38 HE, Colibri2.0 470 light-emitting diode [LED]), yellow fluorescent protein (YFP) (46 HE, Colibri2.0 505 LED) and DAPI (49, HXP 120 C). For cell staining a final concentration of 200uM Hoechst (Hoechst 33342 Solution (20 mM), ThermoScientific) was added and cells were incubated for 10 minutes at 37 degrees. Cells were pelleted by centrifugation at 3,000 × g for 1 min. Supernatant was removed and cells were resuspended in 50 μl of PBS. For single-time-point live-cell imaging, 2 μl of culture was spotted onto a 1.5% low-melting-point agarose pad made in a final concentration of PBS with 0.2% glucose. For pad preparation, agarose was heated to 65°C and poured into a FastWell incubation chamber (Grace Bio-labs, 18×18mm, 1mm depth) pressed onto a 25-mm by 75-mm glass slide. A second slide was pressed down on top, and the setup was allowed to cool at room temperature for 10 min. The top slide was removed, and two microliters of *M. smegmatis* culture was added to the pad then topped with a coverslip (18×18mm, No 1.5).

### Proximity labeling, lysate preparation, and biotinylated protein capture

From a log phase starter culture (OD600 0.3-0.6), 3×30ml cultures for each strain were prepared in 60ml inkwell bottles (Nalgene, #342020-060) targeting a final OD600 of 0.3 after 18hrs (*M. smegmatis*) or 48hs (Mtb). For *M. smegmatis* RpoC-ALFA proximity labeling, the following strains were used: MG7085 (No ALFA tag control) and MGM7106 grown in 30mls LBsmeg with kanamycin 20ug/ml. For Mtb PKS13-ALFA experiment: MGM7416, MGM7418, MGM7419, MGM7420, MGM7421 were grown in 30ml 7H9OADC with kanamycin 20μg/ml. Cultures were washed with 10ml Sauton’s media (biotin-free media) then resuspend 30ml cultures in Sauton’s media with kanamycin in fresh 60ml inkwell bottle and grown at 37C shaking at 150rpm (6h *M. smegmatis*, 24h Mtb) followed by addition of ATc-50ng/ml (3h *M. smegmatis*) or ATc-100ng/ml (18h Mtb). Cells were collected by centrifugation at 3700xG for 10min at room temperature and resuspended in 2ml Sauton’s media (no ATc/no kanamycin) and transferred to 2ml O-ring sealed tube, collected by centrifugation at 3700xG for 5min at room temperature, and resuspended in 1ml Sauton’s media. Cultures were incubated for 2hrs with 200μM Biotin (Sigma, B4501) with shaking at 37C and then collected by centrifugation at 20,000xG for 1min at 4C and placed immediately on ice. Cell pellets were washed with 2ml cold TBS (20mM Tris, 150mM NaCl, pH 7.5), then collected by centrifugation at 20,000xG for 1min at 4C and cell pellets frozen at −20 degrees C.

For lysate preparation, frozen cell pellets were thawed on ice and resuspended with 250ul bead mix (1:1 mix, Biospec 11079101z and 1079107zx) and 1ml cold TBS with protease inhibitors (PI) (Pierce, PIA32955). Lysis was performed by mechanical disruption 3X45 seconds (Biospec Minibeadbeater) with 5 min icing between runs. Unbroken cells/debris were collected by centrifugation at 20,000xG for 1min at 4C, supernatants collected, and residual pellets lysed again with beads and 400ul cold TBS with protease inhibitors. Pooled supernatants were mixed with 150ul of 10x RIPA buffer detergents (TBS+ 10% triton x-100, 1% SDS, 5% sodium deoxycholate) and mixed by inversion, centrifuged at 20,000xG for and (for Mtb) the supernatant was filtered twice through EMD Millipore™ Ultrafree™-CL Centrifugal Filter Devices with Durapore™ Membrane to remove any residual *M. tuberculosis*. For biotinylated protein capture, protein lysates were mixed with 80ul of Streptavidin-Agarose (GoldBio, S-105-10) prewashed 3X in 1ml TBS and incubated rotating at 4C for 18h. Agarose beads were collected at 250xG for 1min at 4C and washed 2xs with 1ml RIPA/SDS (TBS+ 0.1% triton x-100, 0.1% SDS, 0.5% sodium deoxycholate) followed by 4 washes with TBS.

### Peptide preparation for Mass Spectrometry

Streptavidin-agarose beads were resuspended in 40ul of 2M urea, 50mM tris pH 8.0 then 0.4ul of 100mM DTT was added to each sample. Beads were incubated at 37C for 30 min with shaking then 0.3ul 500mM iodoacetamide was added and beads were incubated at room temperature for 45 min protected from light. 0.3ul 500mM DTT was added then 750ng of trypsin protease (Pierce, 90057) and beads were incubated overnight shaking at 37C. Beads were spun down 250xG 2min and supernatant was transferred to a new tube. 500ng of trypsin was added to the supernatant and incubated for 2h at 37C with shaking. Samples were then acidified with trifluoroacetic acid (TFA) to a final concentration of 0.1%. 200ul of 80% acetonitrile (ACN), 0.1% TFA was added to a BioPureSPN Mini C18 spin columns (Nest Group, Product # HUM S18R) and spun at 100 x G for 1 min. 200ul of 0.1% TFA was added to the column and spun 100xG for 1 min, then repeated. The digested sample was then added to the column and spun 100xG for 1 min. 200ul of 0.1% TFA was added to the column and spun 100 x G for 1 min, then repeated. The column was then placed in a fresh tube and peptides were eluted with 100ul 40% ACN, 0.1% TFA and spun 100 x G for 1 min. Peptides were dried in a speedvac.

### Mass spectrometry data acquisition

Samples were analyzed on an Orbitrap Eclipse mass spectrometry system equipped with an Easy nLC 1200 ultra-high pressure liquid chromatography system interfaced via a Nanospray Flex nanoelectrospray source (Thermo Fisher Scientific). Samples were injected onto a fritted fused silica capillary (30 cm × 75 μm inner diameter with a 15 μm tip, CoAnn Technologies) packed with ReprosilPur C18-AQ 1.9 μm particles (Dr. Maisch GmbH). Buffer A consisted of 0.1% Formic Acid (FA), and buffer B consisted of 0.1% FA/80% ACN. Peptides were separated by an organic gradient from 5% to 35% mobile buffer B over 120 min followed by an increase to 100% B over 10 min at a flow rate of 300 nL/min. Analytical columns were equilibrated with 3 μL of buffer A.

To build a spectral library, samples from each set of biological replicates were pooled and acquired in data-dependent manner. Data-dependent acquisition (DDA) was performed by acquiring a full scan over a *m/z* range of 375-1025 in the Orbitrap at 120,000 resolving power (@ 200 *m/z*) with a normalized AGC target of 100%, an RF lens setting of 30%, and an instrument-controlled ion injection time. Dynamic exclusion was set to 30 seconds, with a 10 p.p.m. exclusion width setting. Peptides with charge states 2-6 were selected for MS/MS interrogation using higher energy collisional dissociation (HCD) with a normalized HCD collision energy of 28%, with 3 seconds of MS/MS scans per cycle.

Data-independent analysis (DIA) was performed on all individual samples. A full scan was collected at 60,000 resolving power over a scan range of 390-1010 *m/z*, an instrument controlled AGC target, an RF lens setting of 30%, and an instrument controlled maximum injection time, followed by DIA scans using 8 m/z isolation windows over 400-1000 *m/z* at a normalized HCD collision energy of 28%.

### Mass spectrometry data analysis

The Spectronaut algorithm was used to build spectral libraries from DDA data, identify peptides/proteins, localize phosphorylation sites, and extract intensity information from DIA data (48). DDA data were searched against the *Mycobacterium tuberculosis* or *M. smegmatis* reference proteome sequences in the UniProt database (one protein sequence per gene, downloaded on June 13, 2022). False discovery rates were estimated using a decoy database strategy (49). All data were filtered to achieve a false discovery rate of 0.01 for peptide-spectrum matches, peptide identifications, and protein identifications. Search parameters included a fixed modification for carbamidomethyl cysteine and variable modifications for N-terminal protein acetylation, methionine oxidation, and for phosphoproteomics samples, serine, threonine, and tyrosine phosphorylation. All other search parameters were Biognosys factory defaults.

Statistical analysis of proteomics data was conducted utilizing the MSstats package in R (50). All data were normalized by equalizing median intensities, the summary method was Tukey’s median polish, and the maximum quantile for deciding censored missing values was 0.999.

## Acknowledgements

This work was funded by P30 AI168433 (Tri-I TRAC) and P30 CA08748.

## SI Figures

**Figure S1.**
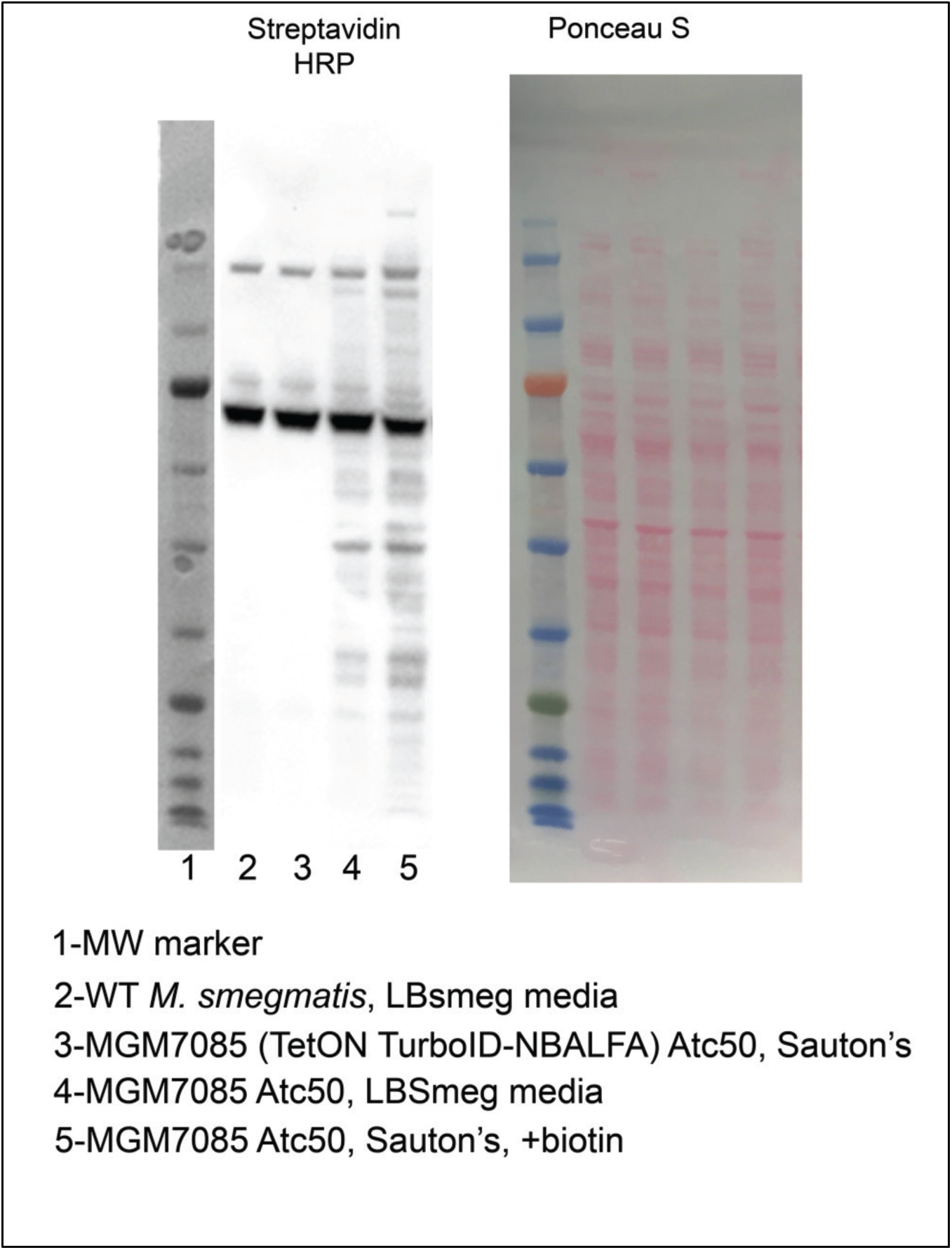
Streptavidin HRP blot (left) and ponceau S staining (right) of the indicated strains and growth conditions showing levels of endogenous biotinylated proteins in *M. smegmatis*.

**Figure S2.**
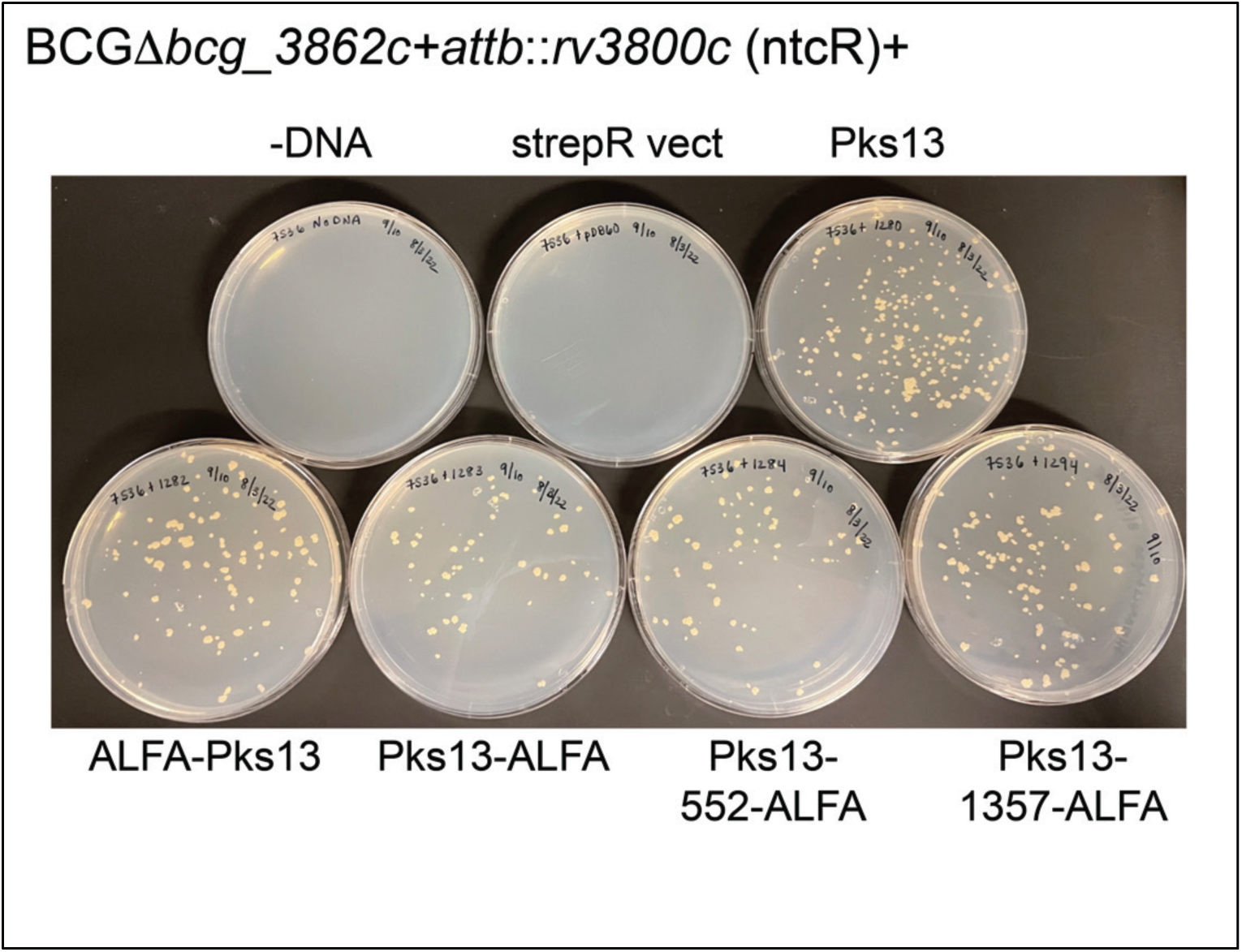
Functionality testing of PKS13-ALFA alleles via *attB* allelic exchange. Shown are streptomycin agar media with transformation of a merodiploid BCG strain with a chromosomal deletion of *pks13* (*BCG_3862c*) and a second copy of M. tuberculosis *rv3800c* at the *attB* site conferring nourseothricin-resistance. Transformation of this strain with an *attB* integrating vector conferring streptomycin-resistance encoding nothing (strepR vect) or alleles of *pks13* with no ALFA tag, or four different ALFA tag position in the PKS13 protein.

**Figure S3.**
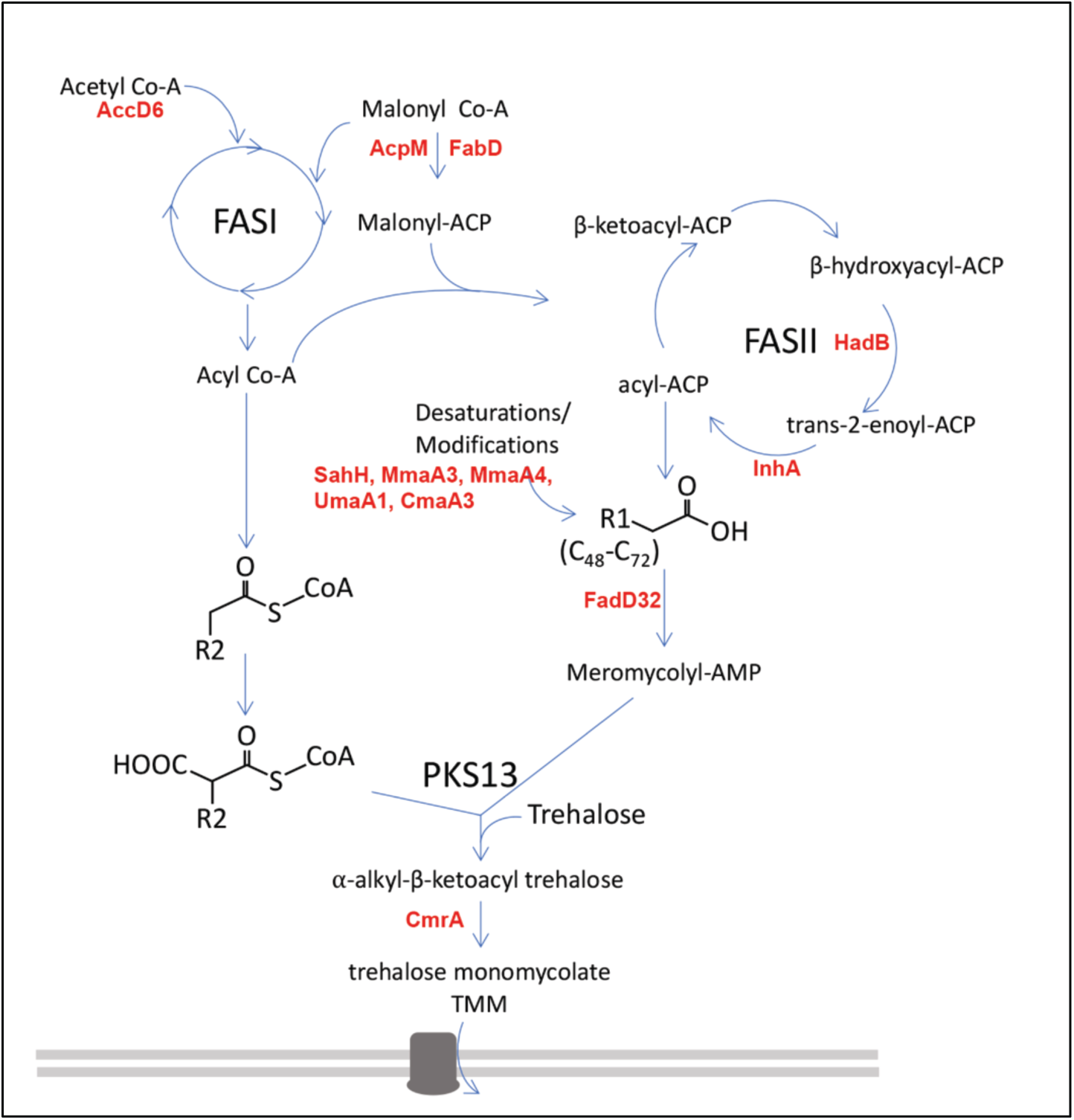
PKS13-ALFA proximity labeled hits within the mycolic acid biosynthesis pathway. Proteins found in all four ALFA tagged labeling conditions with a Log2FC >1.3 and adjusted P-value < 0.05 in bold, red font.

